# Identification of plant-derived microRNAs in human kidney

**DOI:** 10.1101/2022.10.25.513797

**Authors:** Xi Chen, Xishao Xie, Lu Liu, Hongyu Chen, Bo Wang, Zheng Li, Linghui Zeng, Michael P. Timko, Jianghua Chen, Weiqiang Lin, Longjiang Fan

## Abstract

Plant-derived microRNAs (miRNAs) have been implicated as functional regulators in human diseases, although conclusive evidence of this effect remains to be reported. To examine their potential functional role, we profiled the plant-derived miRNAs in 139 blood exosome samples from renal transplantation patients and were able to identify 331 plant-derived miRNAs representing 149 families. According to their miRBase annotation, these miRNAs can be traced back to 76 plant species, most of which are foods common to the human diet (e.g., tomato, soybean, potato and rice). We also profiled 41 blood exosome samples from 22 patients with acute immune rejection (AR) of renal transplants and compared them to 21 samples from 11 patients with stable allograft function to explore possible roles of the functional plant miRNAs. We identified three plant-derived miRNAs (miR4995, miR2118/2218 and miR167) associated with allograft AR whose regulatory targets are mRNAs controlling immune response, T cell activation, and other cellular functions. miR4995 mimics were generated, transfected into HEK293T cells, and their function verified. Our findings not only demonstrate the presence of functional plant-derived miRNAs in human cells, but also provide initial evidence that these miRNAs may be involved in malfunction of renal transplantation.

## 1. Introduction

MicroRNAs (miRNAs) are a class of 20-24 nucleotide small non-coding RNAs that modulate a wide variety of critical biological processes, including cell differentiation, apoptosis, immune responses, and maintenance of cell and tissue identity in animals, plants and other eukaryotes [1,2]. Dysregulation of miRNAs in human cells has been linked to various diseases, including renal allograft status [3,4]. In addition to their occurrence intracellularly, miRNAs have been reported to circulate in biofluids, such as blood [5], urine [6] and saliva [7], and have emerged as potential biomarkers for disease diagnosis and prognosis. Exosomes (extracellular vesicles (EVs)) are the main carriers for circulating miRNAs transferred into neighboring or distant cells in order to modulate cell metabolism [8–10]. Chen et al. [11] recently reported that stomach is a primary site for dietary microRNA absorption and the uptake of these exogenous miRNAs is mediated by SIDT1 expressed on gastric pit cells. Therefore, plant-derived miRNAs could potentially function in cross-kingdom communication. Plant and animal miRNAs have distinct mechanisms of biogenesis and target recognition [12,13]. The loci that produce miRNAs in plants and animals have distinct genomic arrangements, and each kingdom uses different suites of enzymes for miRNA biogenesis [14]. How miRNAs recognize their targets in the two kingdoms have obvious differences [15]. In contrast to the perfect or near-perfect complementarity requirements for targeting of plant miRNAs, the major mode of miRNA target recognition in animals is ‘seed pairing’ at positions of 2 to 8 of the miRNAs [16]. This imperfect complementarity in miRNA∷target interaction gives animals more flexibility in target recognition than plants [17]. However, recent studies have suggested that the line between plant miRNAs and animal miRNAs may not be so clear. Zhang and colleagues [18] were the first to report that plant miRNAs had the potential to function in animal cells in a manner similar to endogenous miRNAs. These researchers reported that a rice miRNA, miR168, is detectable in human serum and demonstrated that miR168 is capable of inhibiting the expression of its target low-density lipoprotein receptor 1 in the liver. It was proposed that rice miR168 was absorbed through the gastrointestinal (GI) tract, packaged into EVs to avoid degradation in the low pH (pH=2) environment of the GI tract, and released into mammalian (human and mouse) liver cells. More recently, Chin and colleagues [19] provided evidence that an ingested plant miRNA, miR159, may exert a therapeutic effect on breast cancer outside the GI tract. miR159 significantly reduced breast tumor growth in mice by targeting a transcription factor *TCF7*. Gadhavi et al. found that miRNAs from *Bacopa monnieri* could target human genes associated with numerous human diseases including cancer [20]. Kalarikkal et al. used edible nanoparticles harboring miRNAs from ginger and grapefruit plants and the expression of the miRNAs in human gastrointestinal tract were confirmed by qRT-PCR [21]. Such observations are not without controversy, since some studies failed to find evidence supporting the presence of plant miRNAs following consumption of various types of food [22–26]. It is possible that the use of robust high-throughput sequencing to screen for all plant miRNAs could resolve the ability to detect these molecules [27].

Acute renal transplant failure, a life-threatening complication, is associated with a high incidence of acute rejection (AR), post-transplantation diabetes mellitus, and delayed grafted function, among other indications [28–30]. Human miRNAs have been reported to be involved in the regulation of the transplantation failure and have been used as biomarkers for preventing the loss of function of transplanted kidney [31–33]. miR-142–5p, miR-155, and miR-223 are overexpressed in AR biopsies [34]. Overexpression of miR-150, miR-155, miR-663, and miR-638 was observed in early cellular rejection biopsies, and four miRNAs were involved in immune response [33]. The possible role of plant-derived miRNA in allograft rejection has not been previously explored and could potentially provide new targets for therapeutic intervention and the development of safer treatment modalities. In the present study, we examined whether plant-derived miRNAs are present in human sera and investigated their potential function as biomarkers for early prediction of or novel therapeutic tools to prevent allograft AR. We show that a wide array of plant-derived miRNAs, representing up to 76 plant species, are detectable in human samples. The presence of three of these plant miRNAs (miR4995, miR2118/2218, miR167) is highly correlated with renal transplant rejection and other renal diseases. In addition, transfection studies with miR4995 mimics further confirmed its involvement in regulating human cellular expression pathways involved in allograft rejection.

## 2. Results

### 2.1. Summary of patients and small RNA profiles

Kidney transplantation is a life-saving treatment modality that is widely used but not without risk or complication. Among the latter is acute rejection that can result in losing allografts in transplant recipients [28,30]. The factors potentiating rejection are not always evident. Therefore, to investigate the potential role of non-endogenous miRNAs in the process, we examined the occurrence of plant-derived miRNAs in renal tissues of normal and AR recipient tissues. A total of 41 samples from 22 AR patients were selected and 21 samples from 11 patients with stable renal allograft function were used as the control group (CG) in this study. The characteristics of included patients were used are summarized in Table S1. Both groups were separated into two subgroups: a preoperative sampling subgroup (e.g., AR-pre, CG-pre) and a postoperative sampling subgroup (e.g., AR-post, CG-post). To examine the presence of plant-derived miRNAs in renal allograft biopsies more extensively, 77 additional blood biopsies from renal allograft patients with various clinical phenotypes were also included for plant miRNA identification. The high-throughput sequencing data were generated from 139 human small RNA libraries with 41 million raw reads on average for each sample. After removing the low-quality tags and contaminants, 27 million clean reads on average per sample were obtained and used in subsequent analyses (Table 1).

**Table 1.**
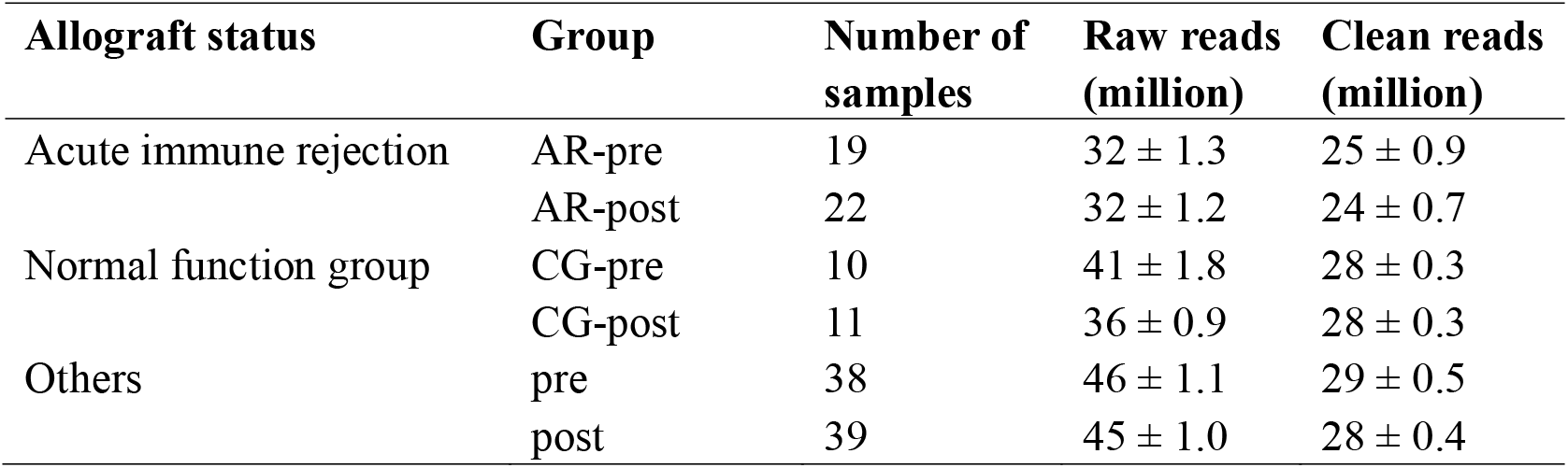
Summary of small RNA sequencing data from 139 blood exosomes of three renal allograft groups by this study. Data are listed as mean ± SEM. AR, acute immune rejection group; CG: normal function group; pre, preoperative sampling subgroups; post, postoperative sampling subgroups

| **Allograft status** | **Group** | **Number of samples** | **Raw reads (million)** | **Clean reads (million)** |
| --- | --- | --- | --- | --- |
| Acute immune rejection | AR-pre | 19 | 32 ± 1.3 | 25 ± 0.9 |
|  | AR-post | 22 | 32 ± 1.2 | 24 ± 0.7 |
| Normal function group | CG-pre | 10 | 41 ± 1.8 | 28 ± 0.3 |
|  | CG-post | 11 | 36 ± 0.9 | 28 ± 0.3 |
| Others | pre | 38 | 46 ± 1.1 | 29 ± 0.5 |
|  | post | 39 | 45 ± 1.0 | 28 ± 0.4 |

### 2.2. Identification and expression of plant-derived miRNAs

Since the accurate identification of plant-derived miRNAs in human blood samples is critical in this study, we utilized a stringent bioinformatics pipeline, including ePmiRNA_finder previously developed by our group to identify plant-derived miRNAs from non-plant small RNA populations in diverse tissues or samples (Figure 1A, details see Methods). Using the pipeline, we identified a total of 331 plant-derived miRNAs belonging to 149 families in the 139 blood exosome samples analyzed (Figure 1B; Table S2). According to their annotation by miRBase V22 (http://www.mirbase.org) [35], the 331 miRNAs could be attributed to 76 plant species, most of which are common foods in the human diet (e.g., tomato, soybean and rice) (Figure S1; Table S2). In order to understand more fully the potential regulatory functions of identified plant miRNAs in renal allografts, the 149 miRNA families were re-classified into 237 seed-specific ones, based on the similarity of ‘seed’ region sequences according to the definition of the seed region of human miRNAs (2 - 8 nt at the 5′ end of the mature miRNA sequence) and re-named as ‘miRNA family_s_serial number’ in this study (Figure 1A; Table S2) (see Materials and Methods). The ‘s’ represents ‘seed-region specific’ and the serial number starts from ‘a’ and is consecutively numbered to distinguish the different seed-region. Compared to plant-derived miRNAs, human miRNAs in these biopsies were much more abundant as expected, with 742 human miRNAs being identified in the 139 samples with a range of 99 to 333 miRNAs per sample (Table S3). The average reads per million (RPM) mapped reads of the human miRNAs were also significantly higher (approximately 100-fold) than those of the plant miRNAs. For example, the average RPM of human miRNAs identified in each AR sample was 8,755 whereas the average RPM of the plant-derived miRNAs in these samples was 164 (Table S3).

**Figure 1.**
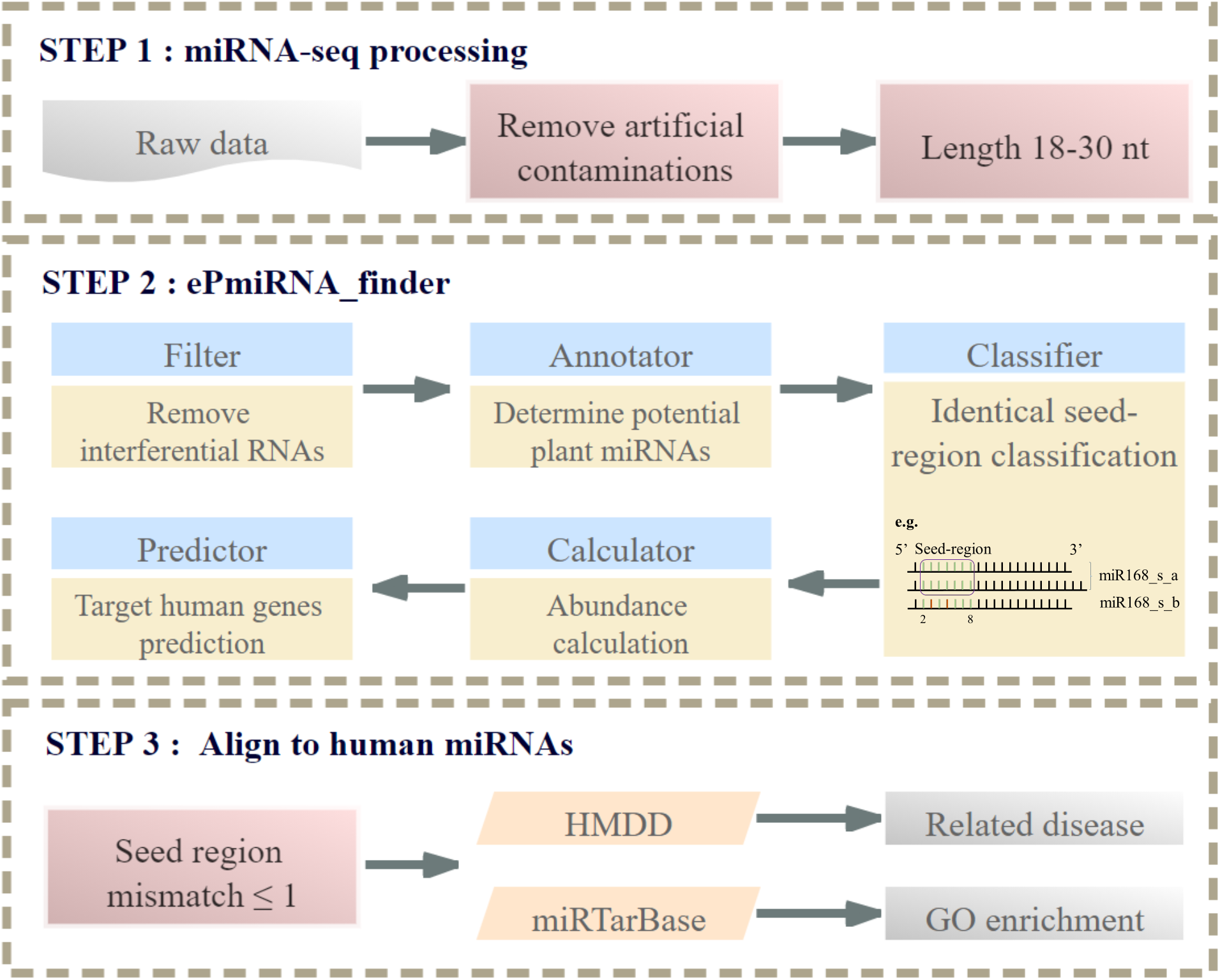
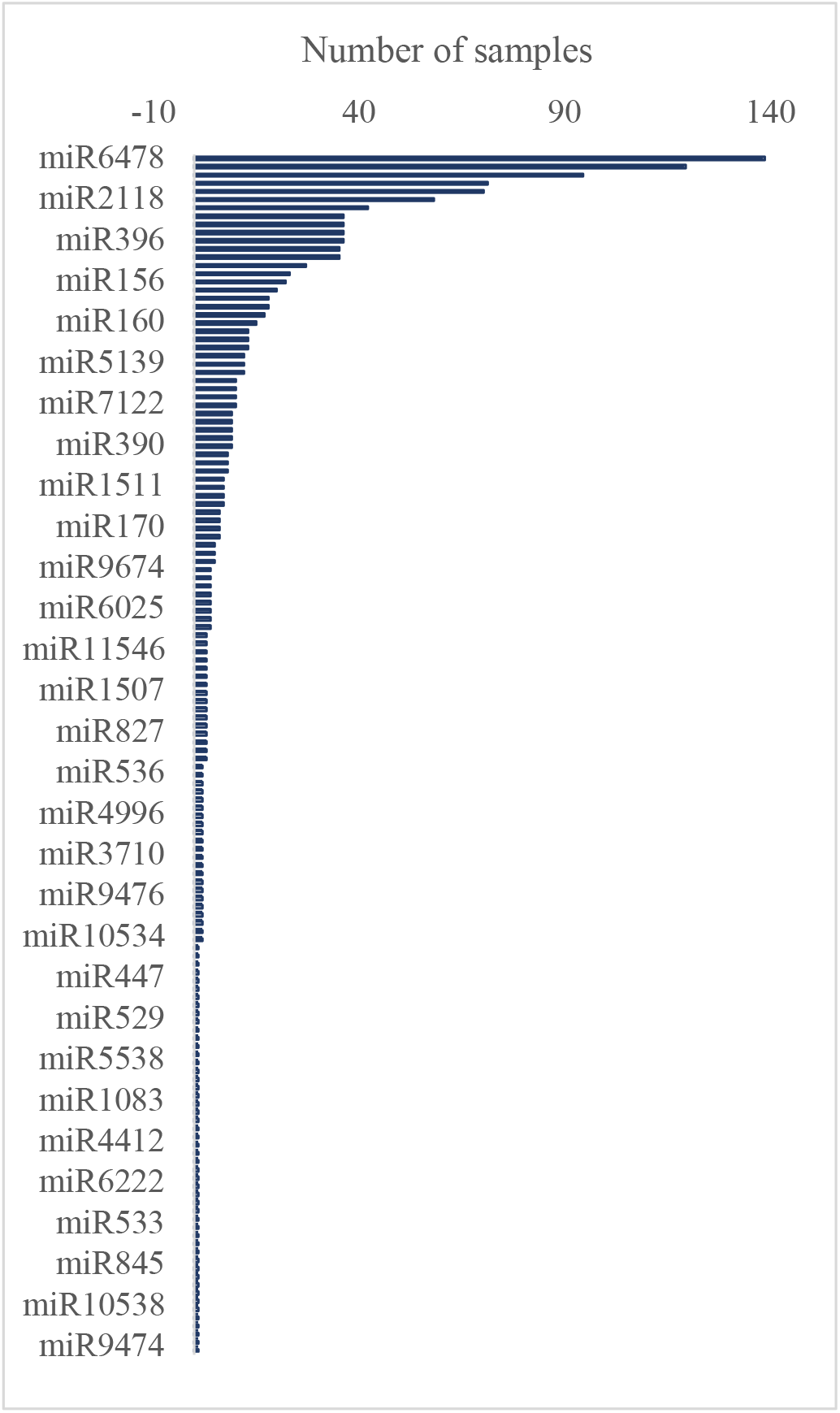
Identification of plant-derived miRNAs in human blood samples. (A) Workflow for identifying plant-derived miRNAs in human miRNA NGS sequencing data and predicting potential human targets. (A) The number of renal allograft samples that plant miRNA families identified.

### 2.3. Plant-derived miRNA distinguishing acute rejection biopsies from normal biopsies

To explore further a potential role of plant-derived miRNAs in acute allograft rejection, we compared the expression profiles of seed-region specific plant miRNAs between AR and CG groups (Figure 2A). On average 5-7 seed-region specific plant miRNAs were detected in at least 30% of the inspected samples in AR or CG (Figure 2A-B). Notable among these was the fact that the plant-derived miR8175 was exclusively present in AR-pre biopsies, while miR4995 was expressed only in CG-pre samples. By comparison, miR168 was missing in AR-pre samples, and there was no significant differential expression in miRNAs between any pairs of samples of the other three groups.

**Figure 2.**
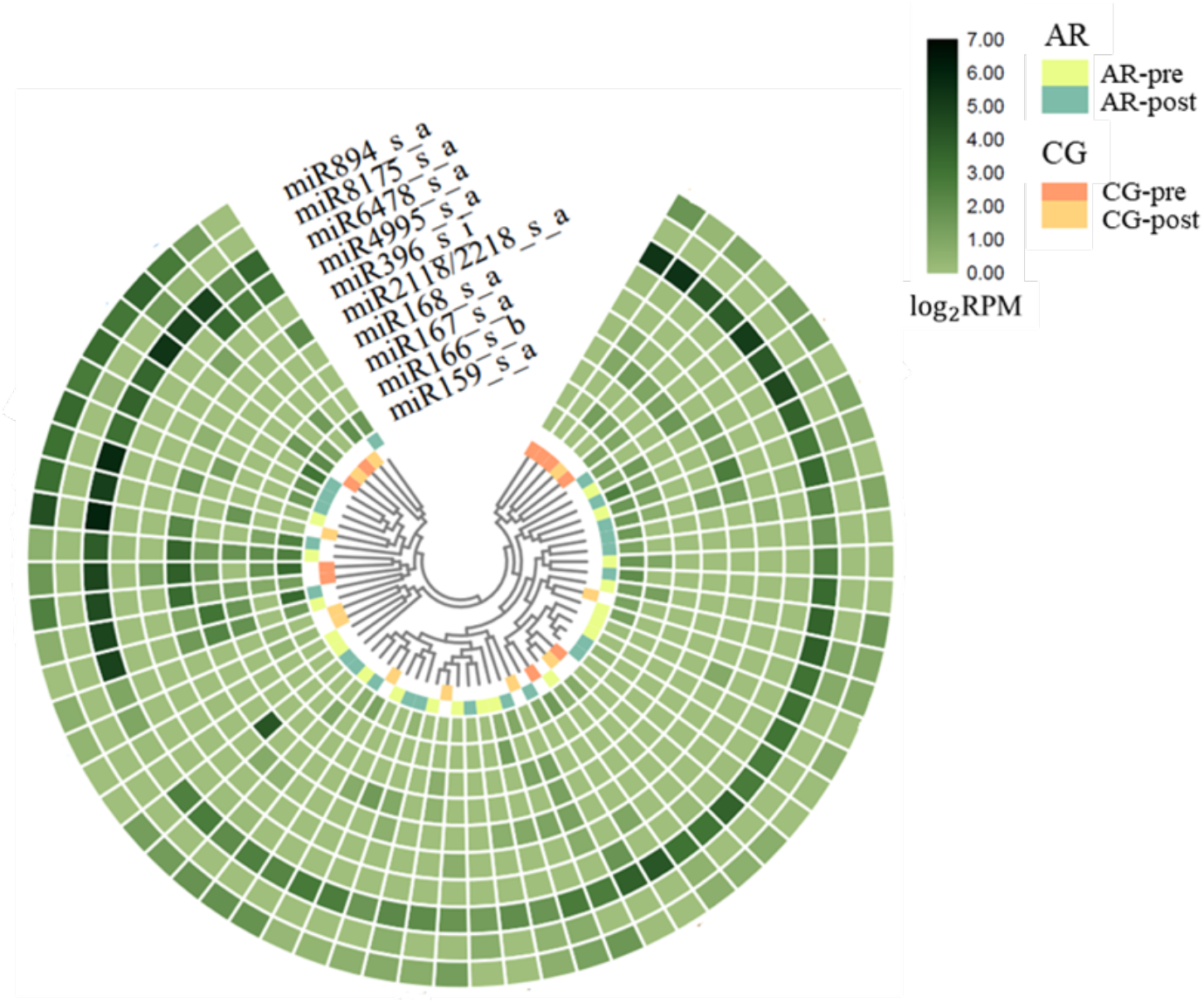
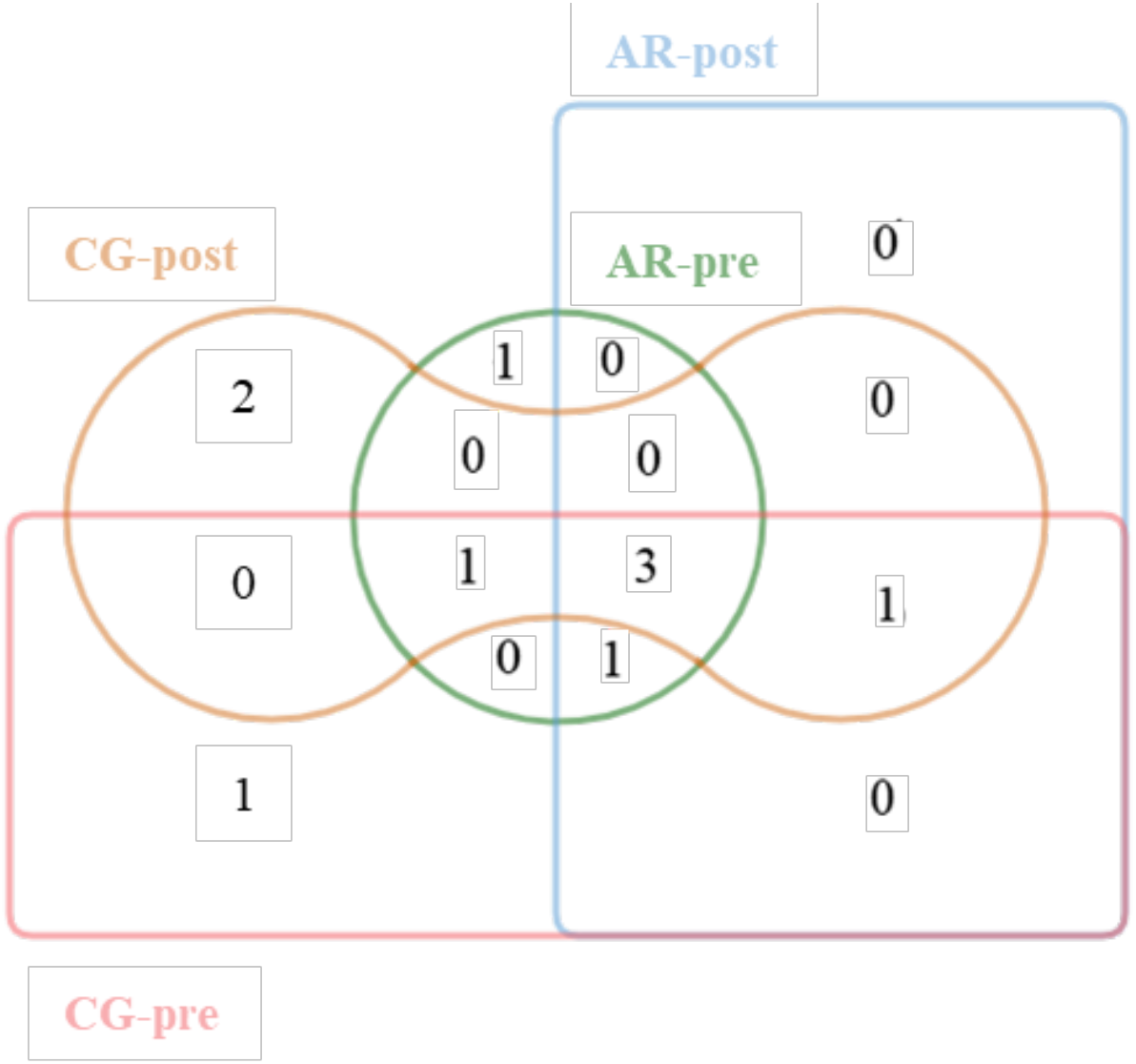
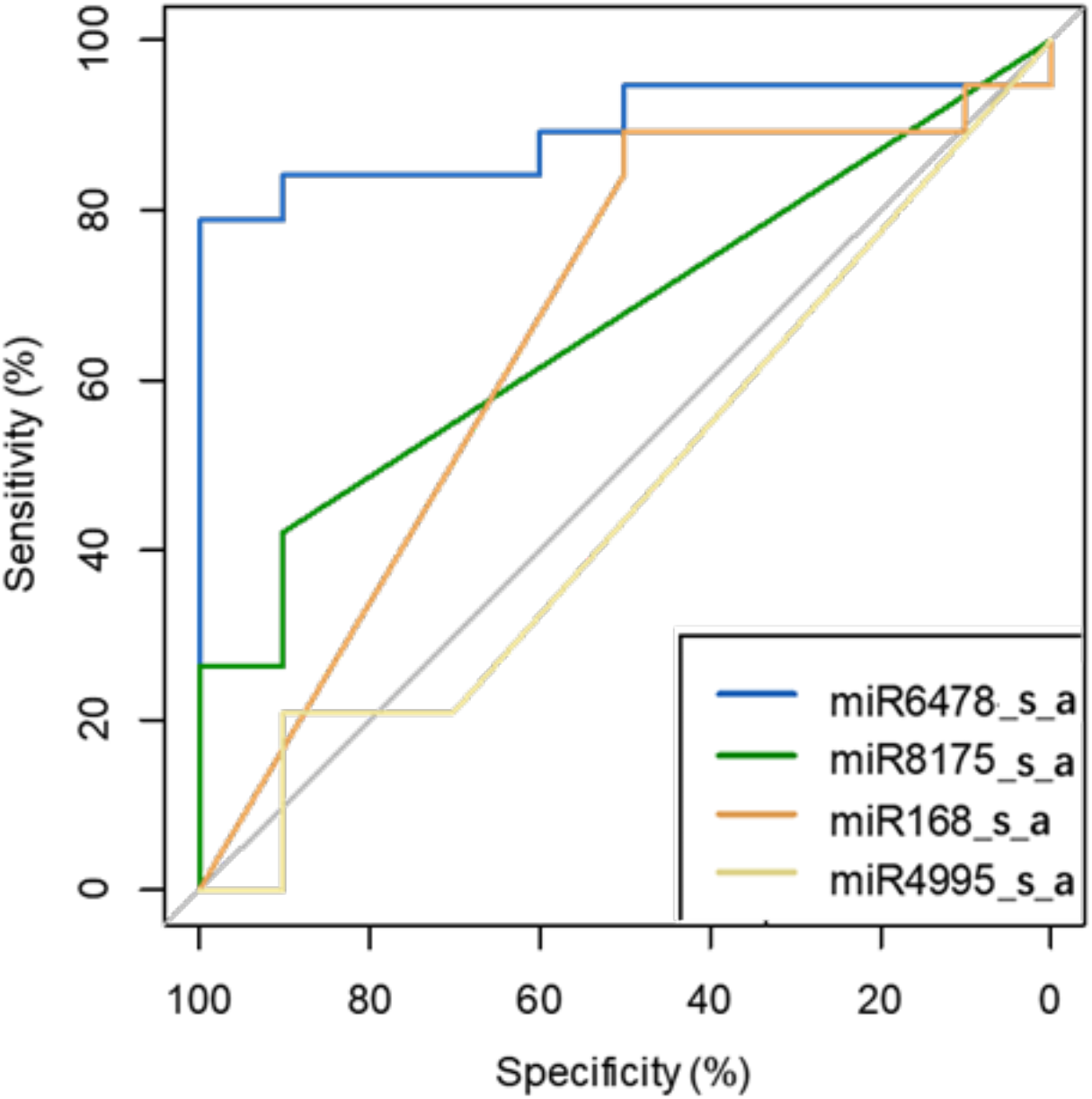
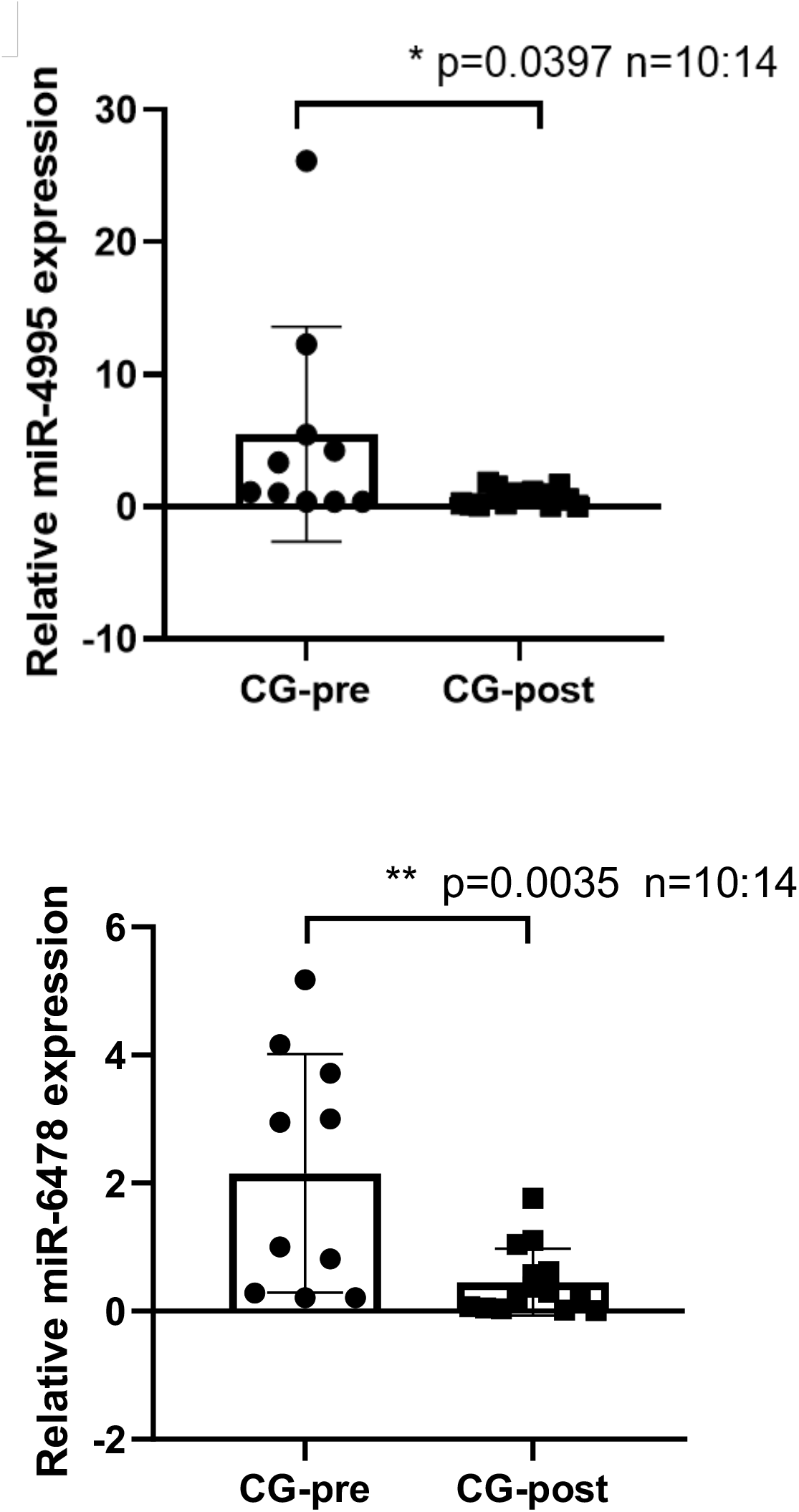
Expression profiles and receiver operating characteristic (ROC) curves of seed-region specific plant miRNAs detectable in at least 30% inspected samples in group AR or group CG. (A) Abundance of seed-region specific plant miRNAs (i.e. with’_s_a’). (B) Distribution of plant miRNAs. (C) ROC curve analyses of four plant miRNAs as potential biomarkers to discriminate between acute immune rejection group and normal function group. (D) Expression level of miR4995 and miR6478_s_a verified by TaqMan RT-PCR. AR: acute immune rejection group; CG: normal function group. Pre: preoperative sampling subgroups; Post: postoperative sampling subgroups.

We also performed receiver operating characteristic (ROC) analysis (a common approach to depict the trade-offs between sensitivity and specificity for a given test [36]), to see whether these three plant miRNAs could be potential biomarkers for prediction of AR. In ROC analysis, the higher the area under the curve (AUC) value, the greater the confidence in the classification effect of that particular biomarker. Analysis of the AUC values of the three seed-region specific miRNAs identified above (i.e., 0.6658 for miR8175_s_a, 0.4658 for miR4995_s_a, 0.6632 for miR168_s_a) did not however convincingly support their role as a potential biomarker (Figure 2C). Seed-region specific miR6478_s_a, which achieved the highest AUC value 0.8947, was ubiquitously found in all selected 62 samples in all four groups. Importantly, miR6478_s_a was expressed significantly higher in CG-pre than any of other three groups (*p* value < 0.05). The optimal threshold point of miR6478_s_a was predicted, and selected through the Youden index. The Yoden index represents the total ability of the classification model to distinguish between rejected and non-rejected patients. The larger the Youden index value, the better performance of the classification model. When the Youden index of miR6478_s_a reached the maximum value of 0.789, the specificity and sensitivity were 1.00 and 0.789, respectively (Figure S2). The optimal threshold value was 11.27, indicating that the possibility of rejection was higher when the expression level of miR6478_s_a was lower than 11.27 RPM. The high values of specificity and sensitivity could suggest that miR6478 is a possible biomarker for rejection. Howevr it needs to be validated in a large clinical sample in the future. Furthermore, we found that seed-region specific plant miR2118/2218_s_a and miR166_s_b were only missing in CG-post and AR-post respectively, while miR167_s_a and miR396_s_i were expressed specifically in CG-post samples. Interestingly, there were no miRNAs found exclusively in group AR or group CG. According to the expression pattern analysis and ROC investigation, all eight plant miRNAs (miR8175, miR4995, miR168, miR6478, miR2118/2218, miR166, miR167, miR396) discovered above have the potential to be involved in the regulation of AR.

### 2.4. Expression validation of plant-derived miRNAs

TaqMan RT-PCR was used to validate the expression levels of miR4995 and miR6478. Both of these plant miRNAs could be detected in blood samples by RT-PCR and their expression trend was the same as that found in the NGS data, with miR4995 and miR6478 being highly expressed in CG-pre, and their abundance being significantly higher than that of other groups (Figure 2D). This not only confirmed the existence of transboundary plant miRNAs, but also further validated the potential of miR6478 as a biomarker.

### 2.5. Functional enrichment of plant-derived miRNA in acute rejection biopsies

Two strategies were applied to determine the potential roles of plant miRNAs in the renal allograft rejection. The first strategy developed based on the assumption that miRNAs with highly similar (< 2 bp mismatch) seed sequences can target to some sets of genes. We aligned the eight plant miRNAs to human miRNAs in target seed regions (Figure S3) and found that the 7-mer sequences of 12 human miRNAs were aligned to miR4995, and 10 human miRNAs could be assigned to miR167, while less than four human miRNAs were aligned to the other six plant miRNAs. We also employed miRTarBase (http://mirtarbase.mbc.nctu.edu.tw/index.html) [37] and HMMD (http://www.cuilab.cn/hmdd) [38] to analyze these sequences. As listed in Table 3, hsa-miR-34a-5p, which is a perfect match for miR4995, was previously reported to be associated with such factors as graft-versus-host disease [39], acute kidney failure [40], lymphocytic B-cell [41], and autoimmune diseases [42]. Interestingly, the experimentally-verified target genes of human miRNAs aligned with miR167 and miR4995 in miRTarBase were enriched into the T cell activation pathway (Figure S4A). It is worth noting that miR2118/2218, which was only aligned to one human miRNA hsa-miR192-3p, could potentially target to *NR1H4* and involved in several kidney injuries, including kidney transplantation rejection [43–45]. For miR168 which previously reported targeting to low-density lipoprotein receptor 1 in the liver, the four matched human miRNAs were associated with autoimmune diseases [46]. For the aligned human miRNAs for plant miR8175, miR6478, miR166 and miR396, there were no records related with kidney or AR in miRTarBase or HMMD.

*De novo* target gene prediction and GO pathway enrichment analyses were then employed as our second strategy. Based on the interaction principle of animal miRNA to target genes [12], we predicted the human target genes for all eight plant miRNAs using ePmiRNA_finder (Table S4). Consistent with the results from the first strategy, the main top GO terms for miR4995 target genes were enriched in immune response and protein kinase B signaling which closed assist immune response in human cells (Figure S4B) [47,48]. The same top terms can be found for miR167 target gene GO pathway enrichment analysis (Figure S4B). Thus, both strategies demonstrated that plant miR4995, miR167 and miR2118/2218 could be involved in allograft rejection by targeting to key human genes.

To verify whether miR4995 influences allograft status by targeting immune response signaling, HEK 293T cells were transfected with miR4995 mimics and control mimics. The transcriptomes of transfected HEK 293T cells were then sequenced using RNA-Seq and differentially-expressed genes (DEGs) were identified. As shown in both KEGG pathway analysis and gene ontology (GO) plots (Figure 3A and 3B), miR4995 targeted genes were enriched in cytokine-cytokine receptor interactions, IL-17 signaling pathway function, inflammatory response, immune system processes, immune response, innate immune response, etc. These results clearly demonstrate the potential regulatory roles of plant-derived miRNAs in allograft rejection.

**Figure 3.**
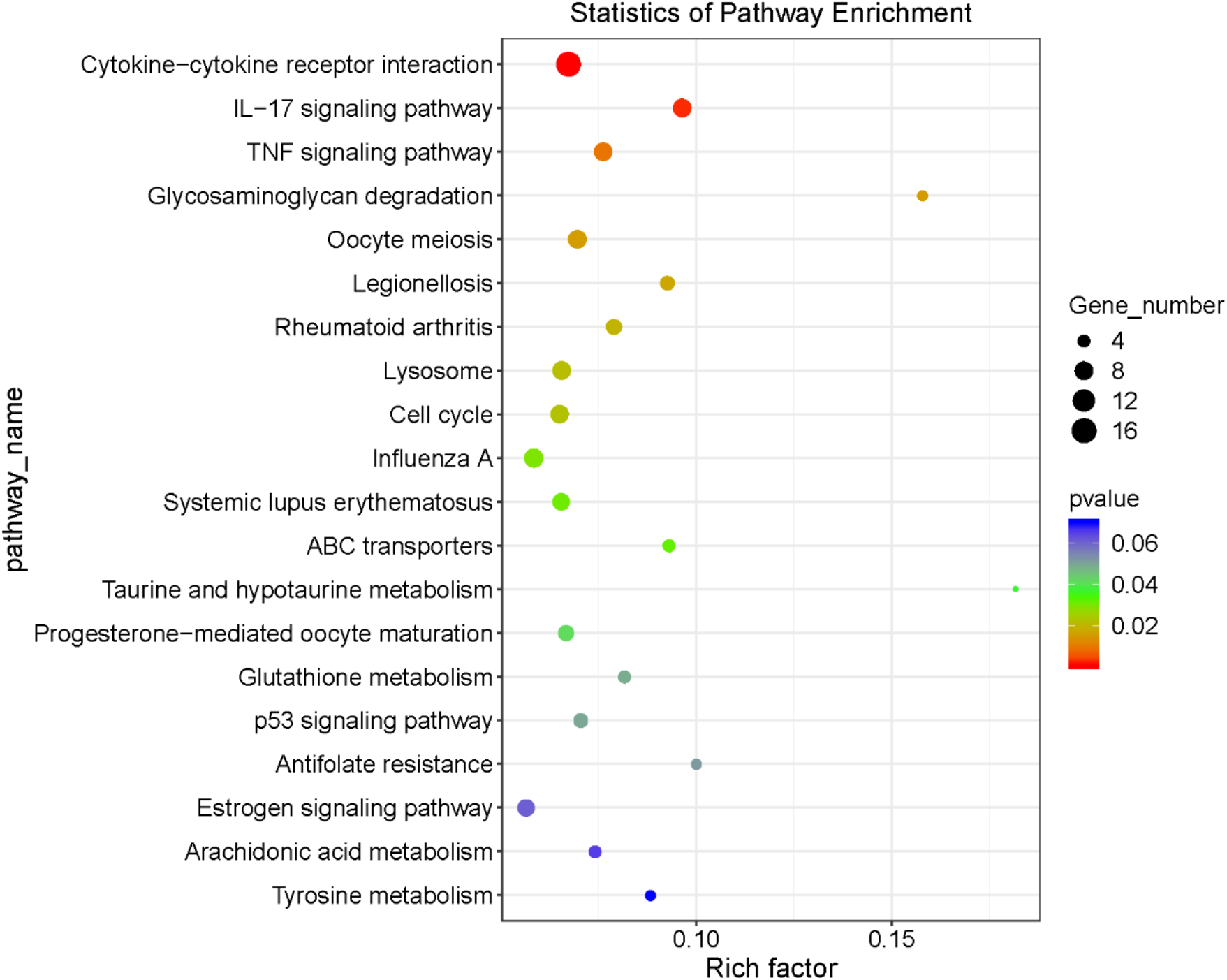
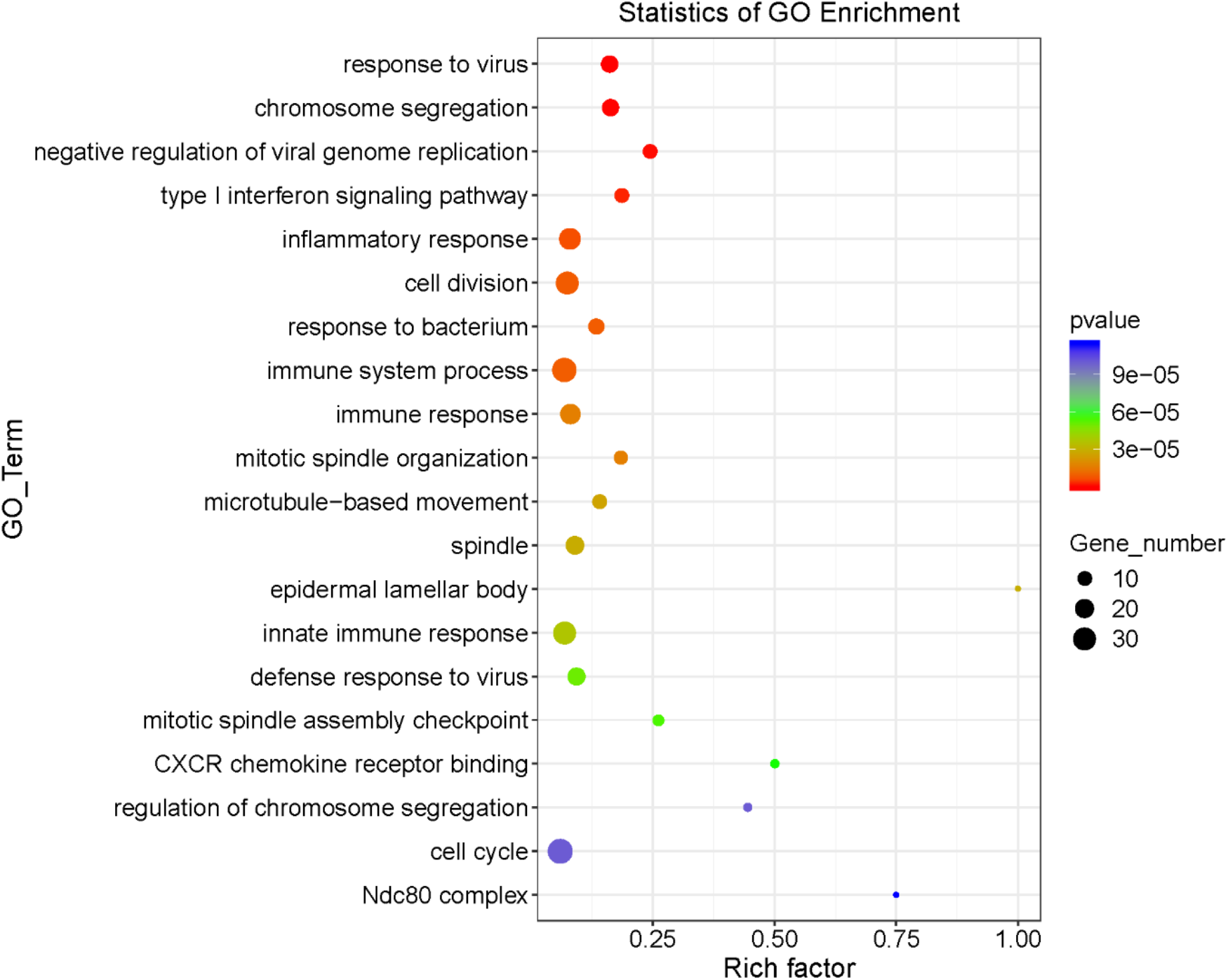
Enrichment analysis of target genes of miR4995 verified by miRNA mimics transfection. (A) KEGG pathways enrichment analysis. (B) GO term enrichment analysis.

## 3. Discussion

There is increasing evidence that exogenous miRNAs could circulate in human cells, tissues, and serum in intact and functional forms supporting the paradigm-shifting hypothesis that plant-derived miRNAs may exert cross-kingdom regulatory effects on human physiological processes [49]. Therefore, plant miRNAs acquired through the human diet could potentially serve as a powerful and affordable treatment for various human maladies or diseases with low toxic side-effects [50]. Chin and colleagues [19] have experimentally demonstrated that plant miR159 significantly reduced breast tumor growth. The present study suggests that plant miRNAs observed in patient’s serum could potentially influence renal allograft status and influence allograft immune rejection. We identified a set of markers associated with which commonly found in the diet and showed that they differ between CG and AR patients. Specifically, miR6478 with ROC value around 0.9 could be used as a biomarker for prediction of AR, and miR4995, miR2118/2218, miR167 could target to human genes and affect the AR pathways (Table 2, Table S4). Therefore, the results of this present study indicate that plant-derived miRNAs could potentially be disease biomarkers, new targets for therapeutic intervention, and assist in the development of safer treatment modalities, even though the candidate miRNAs will subsequently need to be fully validated experimentally with larger clinical population.

**Table 2.**
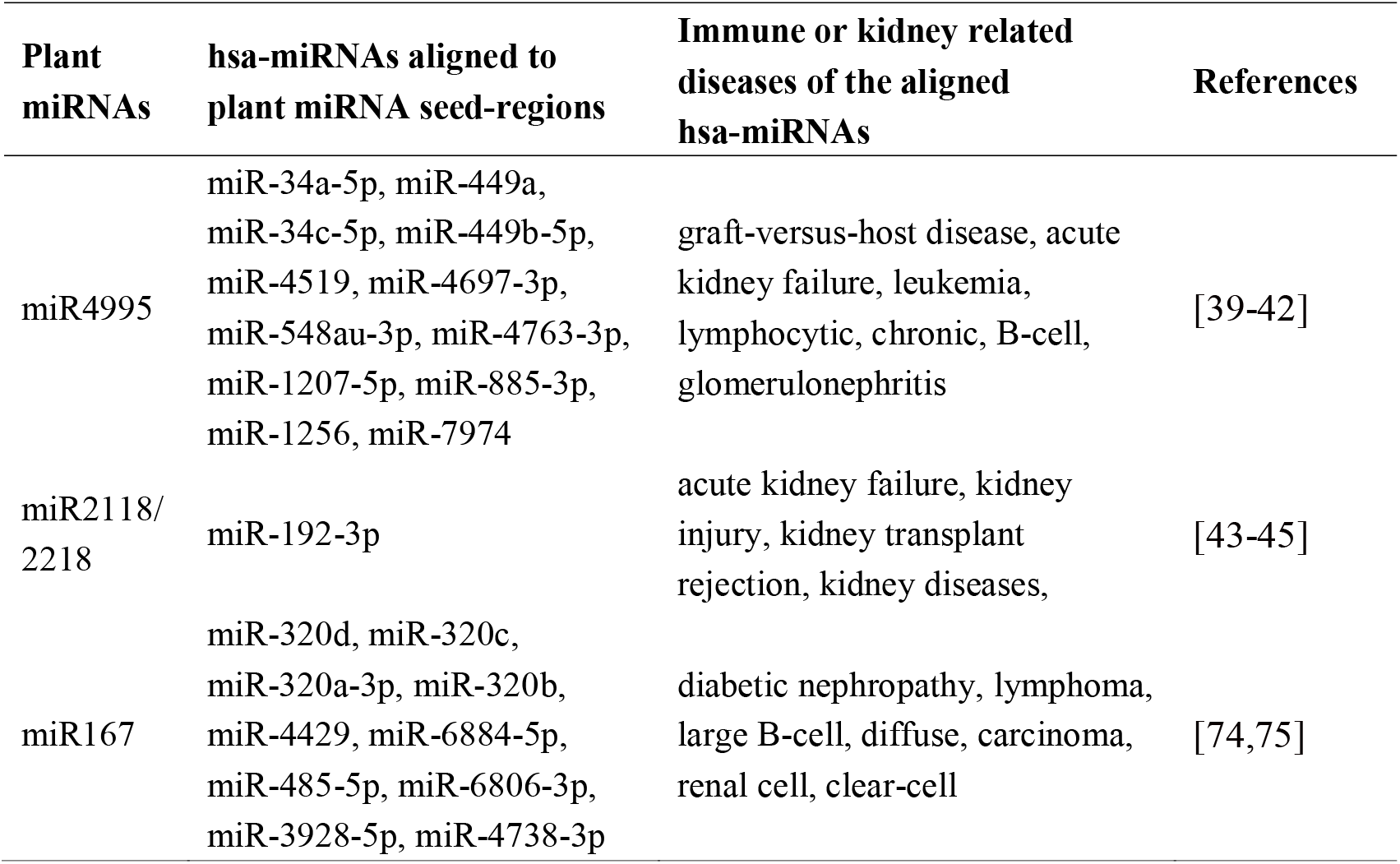
Summary of three candidate plant miRNAs relative to acute immune rejection of renal allograft. Aligned hsa-miRNAs to the candidate plant miRNA seed-regions (i.e. with ‘_s_a’ in Table S3) and related diseases’ references based on the Human microRNA Disease Database (HMDD) were provided.

In present study, we employed multi-layer feature selection methods and classifiers for plant-derived miRNA identification from human miRNA sequencing data. The pipeline (ePmiRNA_finder) has been tested successfully not only in human samples, but also in other species such as pig, cow and mice. It is worth noting that we predicted the targeting of plant miRNAs as human miRNA principle by grouping them according to the ‘seed region’ to improve the prediction accuracy. The result was an ability to identify 331 miRNAs classified into 237 seed-specific ones for differential expression analyses. We also integrated, different strategies to investigate the genes and related diseases targeted by candidate differential expressed seed-specific plant miRNAs, including searching HMDD and miRTarBase, two databases that curate experimentally-supported evidence for human microRNA and disease associations [37,38]. We also de novo predicted the target human genes of candidate seed-specific plant miRNAs and performed enrichment functional analysis to cross validate the potential association with renal allograft rejection.

To understand the potential interaction between plant miRNAs and human miRNAs on regulation of AR of renal allografts, we explored the expression pattern of human miRNAs of group AR and CG. The results showed that the overall human miRNA expression level in AR-pre was the lowest among AR-pre, AR-post, CG-pre and CG-post (Table S5). These data suggest that high expression of specific functional miRNAs could potentially release the allograft rejection stress. Therefore, we investigated the significantly lower expressed miRNAs compared AR-pre to CG-pre, and 17 miRNAs were identified (p value < 0.05, |log2| > 2, min (average RPM in each group) > 1) (Table S5). We further profiled the 8 miRNAs whose average RPM was higher than 100 in CG-pre. We confirmed using the HHDD database that these 8 human miRNAs are also involved in allograft rejection signaling pathways, such as precursor T-cell lymphoma, autoimmune diseases, kidney disease, renal cell carcinoma, and other kidney or IR related diseases (Table S6) [51–55]. Interestingly, there were only 28 overlapping target genes of three candidate plant miRNAs and above 8 human miRNAs (Figure S5), which were enriched in basal pathways that were unrelated to immunity. This suggested that the plant miRNAs and human miRNAs may simultaneously and independently affect the immune rejection status. In total, our data support the hypothesis that plant miRNA miR4995, miR2118/2218 and miR167 could be involve in the regulation of AR after renal transplantation. Among them, miR4995 were further identified by qPCR and functional verified by HEK 293T transfection to show that these miRNAs as being able to target key processes in rejection. It is worth noting that the qPCR was conducted using TaqMan probes for detection and quantitation of plant miRNAs. TaqMan PCR assay has higher specificity and sensitivity over normal qPCR assay, and a commonly used SNP genotyping method in clinical application. The detection system could discriminate a one base-pair mismatch so that the probe can specifically recognize the plant miRNAs. Although these plant miRNAs have been shown to be strongly associated with the allograft AR, further studies are clearly needed.

Several studies have investigated whether small RNAs from plants used in traditional Chinese medicine (TCM) can function as novel precision therapeutic treatments [56,57]. For example, Huang et al. [56] identified thousands of unique small RNAs derived from 10 TCM plants in the blood cells of humans who consumed herbal decoctions and the lung tissue of mice fed these herbal decoctions via oral gavage. Similarly, Du et al. [57] reported that a miRNA from the medicinal herb Hong Jing Tian (HJT, *Rhodiola crenulata*), can effectively reduce the expression of fibrotic hallmark genes and proteins in alveolar cells *in vitro* and in mouse lung tissues I. In traditional Chinese medicine, the HJT has been applied as one of the most effective anti-aging ingredients for centuries. In our study, among all identified plant miRNAs, 46 sequences were perfectly matched to HJT RNA sequences, among which 25 were found in the AR and CG groups (having a total of 4,886 reads). miR159 was also identified in HJT and was present in more than half of the samples, and the expression level also accounted for 1/3 of the total abundance. Based on these findings, our data and those of other laboratories make a strong case for the potential use of dietary plants and those used in TCM as potential starting points for future research on the development of safer and more ‘natural’ treatments of AR.

Overall, we comprehensively mined plant miRNAs from exosome samples of renal transplant patients. We found a small number of plant mature miRNAs differentiated acute rejection from normal ones, and the potentially targeting of plant miRNAs to human genes that strongly positive associated with renal injury, suggest that plant miRNA may serve as biomarkers of allograft status. Our findings add additional evidence to support the observation that plant-derived miRNAs can be found in the human circulatory system and may regulate human gene expression and even impact human diseases such as cancer, influenza, and allograft AR. In addition, diet derived miRNAs from plants could potentially contribute to the early diagnosis or prevention of AR. Clearly, given the controversial nature of these findings, definitive evidence and a full understanding of their role will require additional experimentation.

## 4. Experimental Section

### 4.1. Participants and samples

The Institutional Review Board at the First Affiliated Hospital of Zhejiang University in China approved the study, and all procedures were performed in accordance with institutional guidelines. Informed consent was obtained from all subjects. Blood samples were collected at baseline (the day prior to transplantation) and 2 weeks after transplantation from kidney transplant recipients. The samples were centrifuged at 2000g for 20 min at room temperature, the supernatants collected and stored at −80 ◻ until used. A total of 139 blood samples (5 patients only have the baseline samples) from 72 adult kidney transplant recipients between January 2017 and June 2018 were collected and analyzed. Of the 72 patients, acute rejection occurred in 22 and in all during the first month after transplantation. Patient’s characteristics and laboratory results are presented in Table S1.

### 4.2. Isolation of plasma exosome miRNA

Blood was collected in an EDTA anti-coagulation tube and centrifuged at 4000g for 15 min. The supernatant was collected and centrifuged at 13000g for 10 min, aliquoted, and stored in −80◻ until use. Isolation of exosomal RNA from plasma was performed using the Qiagen ExoRNeasy Serum/Plasma Midi kit (QIAGEN, Cat. 77044) according to the manufacturer’s instructions.

### 4.3. miRNA sequencing data acquisition and processing

High resolution small RNA sequencing of the isolated exosomal RNA was carried out by Beijing Genomics Institute (BGI) using the unique molecular identifiers (UMI) platform. Briefly, enriched and purified small RNAs were used to generate libraries by ligating the 5’-adenylated and 3’-blocked adapter to the 3’-end of the small RNA fragment. After adding the UMI-labeled primer, unligated adaptors were digested and 5’-end adaptor ligation. Single-strand cDNA synthesis was carried out followed by PCR amplification, and the library fragments in the range of 100-120 bp were selected. Libraries meeting quality standards were then sequenced on BGISEQ-500. Adapters, primers and poly A were removed from the raw data (Figure 1A). Only reads with length between 18 nt and 30 nt were kept for further analysis. Then, all low-quality tags were eliminated from the datasets, which have more than four bases and whose quality been less than ten, or which have more than six bases and have a quality less than thirteen.

### 4.4. Human miRNA identification

Human miRNA identification was performed according to previously published identification methods [58]. Briefly, quantification of known miRNAs was done on the basis of miRBase precursors and corresponding mature miRNAs. Bowtie 1.1.2 [59] was used to map clean reads to the *Homo sapiens* reference genome (hg38, from http://hgdownload.soe.ucsc.edu/goldenPath/hg38/bigZips/hg38.fa.gz) [60]. Human miRNAs were identified and quantified by miRDeep2 [61] using mapped reads.

### 4.5. Plant-derived miRNAs identification

Plant-derived miRNAs identification was mainly performed by ePmiRNA_finder, which was previously developed by our group for plant-derived miRNA prediction from non-plant small RNA populations in diverse tissues or samples [62]. Briefly, in the first step (filter), the clean reads were aligned to human miRNA/rRNA/tRNA/snoRNA/snRNA/piRNA (available at miRBase 22.1 release [35], Rfam 14.1 release [63], respectively) allowing two mismatches and the unaligned reads were aligned to *H. sapiens* genomes (hg38) allowing one mismatch (Figure 1A). The remaining reads were further compared to sequences from all human associated microbiomes (Human Microbiome Project, http://www.hmpdacc.org/HMRGD/) [64] allowing one mismatch as recent studies have demonstrated that bacteria, fungi and archaea are abundant in human plasma [65]. In the second step (annotator), the unaligned reads were aligned to plant miRNAs using the following four criteria: (1) A maximum of one mismatch is allowed; (2) No `N’ bases in the reads; (3) Sequence coverage that differed by the aligned plant mature miRNA no more than one nucleotide; (4) Each read is assigned to only one optimal plant miRNA. In the third step (classifier), the candidate plant miRNAs from the Annotator module were sorted based on ‘seed region’ matching. Seed region denotes the sequence of 2 - 8 nt at the 5′ end of plant mature miRNA sequences. The classification based on ‘seed region’ of all miRNAs of all plant species in miRBase 22.1 has already been performed and used as reference in this step. The classifications follow the following rules: miRNAs from the same family with identical seed region were merged into one subgroup, termed as seed-region specific miRNA and named as ‘miRNA family_s_serial number’. The ‘s’ represents ‘seed-region specific’ and the serial number started from ‘a’ and is consecutively numbered to distinguish the different seed-region. In this step, the candidate plant-derived miRNAs are classified according to the reference and the counts are re-calculated. The abundance of plant seed-region specific miRNAs was normalized by modified RPM:

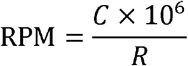

where *C* denotes the count of plant miRNAs, and *R* represents the count of clean mapped reads.

### 4.6. Potential human targets and related diseases predictions

Two strategies were applied to identify potential human targets of plant miRNAs. The first strategy is based on the theory that target genes of human miRNAs with close target seed sequences were assumed to be potentially targeted by the plant miRNAs. Seed region of the plant miRNAs were extracted and aligned with the seed region of human miRNAs, less than or equal to one mismatch. These matched human miRNAs were searched in HMDD [38] and miRTarBase [37] for the diseases and target genes that had been verified by experiments. Target genes were enriched in GO terms by clusterprofile [66].

The second strategy directly predicts the plant miRNAs target genes using the predictor module of ePmiRNA_finder (https://github.com/bioinplant/ePmiRNA_finder). All human target genes based on seed region of the candidate plant miRNAs were predicted. Four main human miRNA∷target prediction tools were integrated including miRanda [67], RNAhybrid [68], TargetScan [69] and PITA [70]. The exact miRNA binding sites predicted by at least 3 out of the 4 prediction algorithms are chose as targets of candidate seed-specific plant miRNAs. The *H. sapiens* 3′ UTR sequences, that served as potential plant miRNAs targets, were downloaded from the UCSC Bioinformatics Site (http://genome.ucsc.edu/) [71]. The disease-related gene set from the REACTOME database (http://www.reactome.org/) was collected [72], and the target genes that overlap with the gene set were used. GO annotation was performed with these selected genes.

### 4.7. Statistical analysis

ROC analysis was conducted using the ‘pROC’ package in R [73]. Continuous data were reported as mean (± SEM) and analyzed by unpaired Student’s *t* test. Significance levels were pre-specified as having a *p*-value = 0.05.

### 4.8. Quantitative RT-PCR

Reverse transcription and quantitative PCR were performed according to manufacturer’s instruction (Hairpin-itTM microRNA and U6 snRNA Normalization RT-PCR Quantitation TaqMan® Kit, GenePharma). PCR cycle conditions are as follows: 95°C for 3 minutes, followed by 40 cycles of 95°C 12s and 62°C 40s. The primers for microRNA are listed in Table S7. The relative quantities of selected miRNAs were normalized to that of U6.

### 4.9. Cell culture, miRNA mimics transfection

Human embryonic kidney cell line (HEK 293T) was purchased from the American Type Culture Collection and cultured in DMEM medium (GIBCO, Cat. NO. 12430) supplemented with 10% FBS (GIBCO, Cat. NO. 10099) and 1% penicillin/streptomycin (GIBCO, Cat. NO. 15140). The cells were maintained in a 37◻°C incubator with 5% CO_2_. For miRNA transfection, 293T cells were plated on 6-wells culture plate. At the next day, miR4995 mimics or control mimics (purchased from Genepharma) was transfected into 293T cells, respectively, using Lipofectamine 3000™ Transfection Reagent (Invitrogen, Cat. NO. L3000015). Each mimics transfection were conducted with three replicates. Two days after transfection, the cells were collected, suspended with TRIzol reagent. Total RNA was extracted and transfection efficiency was confirmed using Taqman qPCR.

### 4.10. RNA Sequencing and Data Analysis

In order to find out the miRNA targets and related signaling pathway, total RNA was subjected to RNA sequencing (RNA-Seq). The sequencing data were analyzed at omicstudio platform (www.omicstudio.cn). The differential expressed gene (DEG), KEGG and GO analyses were conducted.

## Acknowledgments

This work was supported by the National Key R&D Program of China (2018YFC2000400), and National Natural Science Foundation of China (81670651, 81970573, 81770752, and 81970651).

## Author contributions

L.F., W.L. and J.C. conceived, designed, and supervised the study. X.C. and L.L. wrote the paper. W.L. carried out the validation experiments. X.X., B.W., H.C. and Z.L. collected and analyzed the data. M.P.T. and L.Z. discussed the data and assisted in writing the paper. All authors reviewed and approved the final manuscript.

## Supplementary files

**Supplementary figures:**

**Figure S1.** The number of plant miRNA families identified by this study in potential sources of plant species.

**Figure S2.** The optimal threshold point of ROC curve of miR6478_s_a.

**Figure S3.** The seed region sequence alignments of eight specifically expressed seed-region specific plant miRNAs (i.e. with ‘ _s_a ‘) to human miRNAs. (A) miR8175_s_a; (B) miR4995_s_a; (C) miR6478_s_a; (D) miR168_s_a; (E) miR2118/2218_s_a; (F) miR166_s_b; (G) miR167_s_a; (H) miR396_s_i. * indicated the same nucleotide.

**Figure S4.** GO pathways enrichment analysis of predicted target genes by plant miR4995 and miR167. (A) Target genes of human miRNAs aligned to miR4995 or miR167 provided by miRTarBase. (B) Target genes of miR4995 or miR167 predicted by ePmiRNA_finder. Y-axis represents biological process pathways, and x-axis represents plant miRNAs. Size and color of the bubble represents the amount of genes enriched in the pathway and enrichment significance, respectively.

**Figure S5**. Target genes of eight highly expressed human miRNAs and three candidate plant miRNAs relative to acute immune rejection of renal allograft. Red nodes represented human miRNAs, mazarine nodes represented plant miRNAs, yellow nodes represented genes that could be targeted by plant miRNAs, wathet nodes represented genes that could only be targeted by human miRNAs. The connection indicated targeting relationship.

### Supplementary tables

**Table S1.** Summary and characteristics of renal transplantation patient population used by this study

**Table S2.** Information of plant-derived miRNAs identified from small RNA NGS data from blood exosomes of five renal allograft groups by this study.

**Table S3.** Abundance of human miRNAs identified from small RNA NGS data from blood exosomes of five renal allograft groups by this study.

**Table S4.** Summary of target genes of eight specifically expressed seed-region specific plant miRNAs provided by PmiRNA_finder and miRTarBase.

**Table S5.** Summary of significantly differentially expressed human miRNAs between group A and group C. Group A: acute immune rejection group; group C: normal function group sampling one month after surgery.

**Table S6.** Information of target genes provided by PmiRNA_finder, miRTarBase and HMDD of eight highly expressed human miRNAs relative to acute immune rejection of renal allograft.

**Table S7.** Primers used in RT-PCR for miRNA.

